# Increasing the sensitivity, recovery, and integrality of spatially resolved proteomics by LCM-MTA

**DOI:** 10.1101/2022.08.21.504675

**Authors:** Lei Gu, Xumiao Li, Ziyi Li, Qinqin Wang, Kuo Zheng, Guanyu Yu, Chaoqun Dai, Jingquan Li, Benpeng Zhao, Huiping Zhang, Qizhi He, Wei Zhang, Chen Li, Hui Wang

**Author notes:** **Correspondence** Qizhi He, Wei Zhang, Chen Li, Hui Wang. These authors contributed equally to this work (Lei Gu, Xumiao Li, Ziyi Li).

## Abstract

Conventional proteomic approaches neglect tissue heterogeneity and spatial localization information. Laser capture microdissection (LCM) can isolate specific cell populations or histological areas from heterogeneous tissue specimens while preserving spatial localization information. Formalin-fixed paraffin-embedded (FFPE) is currently a standardized method for long-term stable preservation of clinical tissue specimens. However, spatially resolved proteomics (SRP) studies of FFPE tissues by combined LCM and mass spectrometry (MS)-based proteomics face challenges, such as formalin-induced protein crosslinking limits protein extraction and digestion, protein loss during sample preparation, and the detectability of MS for trace tissues. Therefore, it is necessary to specifically develop SRP sample preparation methods and MS methods suitable for trace FFPE tissues. Here, we provide an SRP method suitable for trace FFPE tissues produced by LCM, termed LCM-Magnetic Trace Analysis (LCM-MTA), which can significantly increase the sensitivity, recovery, and integrality of SRP. The starting material has been reduced to about 15 cells, which resolution is comparable to existing spatially resolved transcriptome (SRT). We also apply our LCM-MTA into SRP studies on clinical colorectal cancer (CRC) tissues and accurately distinguish the functional differences of different cell types. In conclusion, LCM-MTA is a convenient, universal, and scalable method for SRP of trace FFPE tissues, which can be widely used in clinical and non-clinical research fields.

## Introduction

Proteins are actual executors of cellular functions. Recent advances in mass spectrometry (MS)-based proteomic technology have facilitated proteomic studies on clinical tissue samples, thereby promoting the understanding and treatment of diseases^1,2^. However, conventional proteomic methods generally treat clinical tissues as bulk samples for protein extraction, enzymatic digestion, and analysis, resulting in neglecting cellular heterogeneity and loss of spatial localization information^3^. Understanding the expression levels and spatial distribution of genes and proteins during disease initiation and progression is essential for in-depth study of disease biological processes and treatments^4,5^. Spatially resolved transcriptomics has been applied in several biomedical research fields^6-8^ and has been selected as the Method of the Year 2020 by Nature Methods^9^, whereas the development of spatially resolved proteomics (SRP) has failed to meet our needs and is one of the bottlenecks in the field of proteomics. Laser capture microdissection (LCM) can isolate specific cell populations or histological regions from heterogeneous tissue specimens while preserving spatial localization information, and its development makes it possible to specifically perform SRP studies on one or a type of cell in tissue samples^10^. Both fresh-frozen tissues and Formalin-fixed paraffin-embedded (FFPE) tissues are generally common methods for preserving patient tissue specimens in clinical studies. Based on fresh-frozen tissues or FFPE tissues, it provides a valuable resource for clinical SRP research by combining LCM and MS-based proteomics.

Fresh frozen tissue, which can maintain biomolecules in their native state, is considered to be one of the first choice for clinical omics studies. Xu et al^11^. reported an SRP by combing LCM with integrated proteomics sample preparation technology SISPROT, and they identified ∼500 proteins from 0.1 mm^2^ and 10 μm thickness fresh-frozen colon cancer tissue sections. Zhu et al^12^. reported a method to SRP by coupling LCM with Nanodroplet Processing in One pot for Trace Samples (nanoPOTS). And they can identify 180, 695, or 1,827 protein groups from 12-μm-thick fresh-frozen rat brain cortex tissue sections having diameters of 50, 100, or 200 μm, respectively. Davis et al^13^. also reported a method for spatial, Cell-Type-Resolved proteomics of the human brain, and they identified ∼1,500 proteins from 0.06 mm^2^ and 10 μm thick fresh-frozen human brain tissues. However, fresh-frozen tissue degrades rapidly at room temperature and is expensive to store, which is not conducive to long-term preservation. FFPE is currently a standardized method for long-term stable preservation of clinical tissue specimens, which is fixed with formaldehyde solution through cross-linking between proteins and then embedded in paraffin blocks to facilitate processing and long-term preservation^14,15^. LCM can be used to obtain regions of interest or specific types of cells from FFPE tissues, which can provide a valuable resource for SRP research. Griesser et al^16^. quantified over 5,600 protein groups from human substantia nigra FFPE tissue (∼ 3,000 cells) by combining LCM and the single-pot, solid-phase-enhanced sample-preparation (SP3) proteomic sample preparation methods. Buczak et al.^17^ reported an SRP analysis protocol using LCM and SP3 for hepatocellular carcinoma (HCC) FFPE tissues, and they can quantify close to 6,000 protein groups using label-based tandem mass tags (TMT) approach from ∼ 40 mm^2^×10 μm tissues. As a matter of fact, formalin-induced protein crosslinking limits protein extraction efficiency and digestion into peptides, and introduces many non-specific and irreversible protein modifications, thus affecting protein identification, especially for trace tissues containing low-level proteins^18^. Sigdel et al^19^. reported a Near-Single-Cell proteomics approach, but the amount of proteins extracted from FFPE tissue was significantly less than that from fresh-frozen kidney tissue. Herrera et al.^20^ also reported a spatially resolved micro-proteomic technique for human lung FFPE tissues, and they identified more than 1,000 protein groups from 0.0125 mm^3^ (2.5 mm^2^ × 5 μm) tissue. However, excessive target tissue area can drastically reduce spatial resolution, resulting in information homogenization in particular tissues or cell types. Moreover, to improve spatial resolution, LCM samples typically contain only a few dozen cells and contain proteins as low as sub-microgram levels. However, proteins at the sub-microgram level are easily lost during sample preparation and require highly sensitive MS for detection. Therefore, in order to obtain high-resolution SRP information from trace FFPE tissues by combining LCM and MS technology, it is necessary to specifically develop SRP sample preparation methods and MS methods suitable for trace FFPE tissues.

Here, we provide an SRP method suitable for trace FFPE tissues produced by LCM, termed LCM-Magnetic Trace Analysis (LCM-MTA), which can significantly increase the sensitivity, recovery, and integrality of SRP. LCM-MTA can improve the depth of protein identification in trace FFPE tissues and reduce the starting material to about 15 cells, which resolution is comparable to existing spatially resolved transcriptome (SRT). In addition, we verified the stability, universality, and extensibility of LCM-MTA under different staining methods, different section thicknesses, different LCM instrument, and different MS data acquisition modes. We also demonstrated that LCM-MTA can be applied to SRP studies on clinical colorectal cancer (CRC) FFPE tissues, which gives great confidence in our application of LCM-MTA in a wide range of studies.

## Results

The LCM-MTA workflow consists of five major stages shown in Figure 1. Briefly, trace samples of interest were obtained from FFPE tissue using LCM and collected on adhesive caps. Next, trace target tissues on adhesive caps were exposed to lysis buffer and subjected to lyse the tissues in situ. Subsequently, the lysate was centrifuged to collect proteins. In the next stage, reducing agents and alkylating agents were added to the solution. And then beads and trypsin were added to the solution to digest proteins. Finally, digested peptides can be used for quantitative LC-MS/MS analysis.

**Figure 1.**
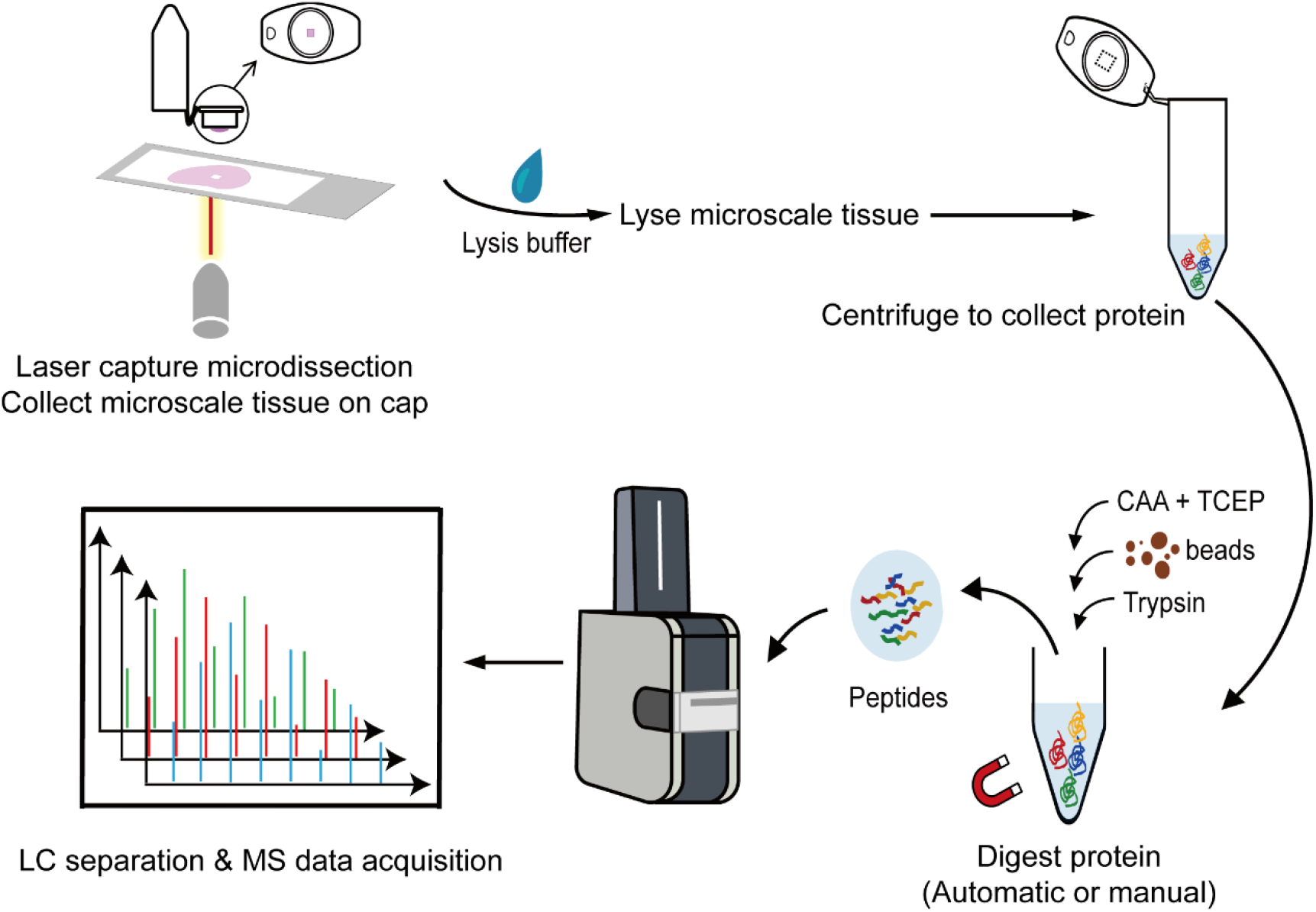
The workflow for spatially resolved proteomic analysis of trace tissues by LCM-MTA. This workflow consists of five major stages. Briefly, trace samples of interest were obtained from FFPE tissue using LCM and collected on adhesive caps. Next, trace target tissues on adhesive caps were exposed to lysis buffer and subjected to lyse the tissues. Subsequently, the lysate was centrifuged to collect proteins. In the next stage, reducing agents and alkylating agents were added to the solution. And then beads and trypsin were added to the solution to digest proteins. Finally, digested peptides can be used for quantitative LC-MS/MS analysis.

### LCM-MTA can improve the depth of protein identification in trace FFPE tissues and reduce the starting tissues to 15 cells

First, we compared the protein recovery of LCM-MTA lysis buffer and conventional sodium dodecyl sulfate (SDS) lysis buffer. For equal amounts of starting tissue (human placenta FFPE tissue, ∼1 mm^2^ × 8 μm), LCM-MTA lysis buffer yielded more protein than conventional SDS lysis buffer based on tryptophan-based fluorescence quantification protein concentration (Figure 2A) and the number of proteins identified by MS (Figure 2B). We also compared the protein recovery of different lysis modalities. Compared with conventional lysis, lysis in situ of LCM-MTA identified more proteins for equal amounts of starting tissue (∼0.1 mm^2^×8 μm) (Figure 2C). All of these indicate that LCM-MTA has a higher protein recovery. In addition, we obtained different areas of human placenta FFPE tissue (section thickness of 8 μm) by LCM (Figure 2D) and trace target tissues were subjected to LCM-MTA and MS. The number of proteins identified in trace tissues of different areas is shown in Figure 2E. Surprisingly, based on MS data-dependent acquisition (DDA) mode, more than 500 proteins can still be quantified at trace tissue size as small as 0.00004 mm^3^ (∼15 cells, 0.005 mm^2^ × 8 μm), which resolution is comparable to existing SRT^5,21^. It also suggests that LCM-MTA can significantly reduce the starting tissues required for individual samples and improve spatial resolution.

**Figure 2.**
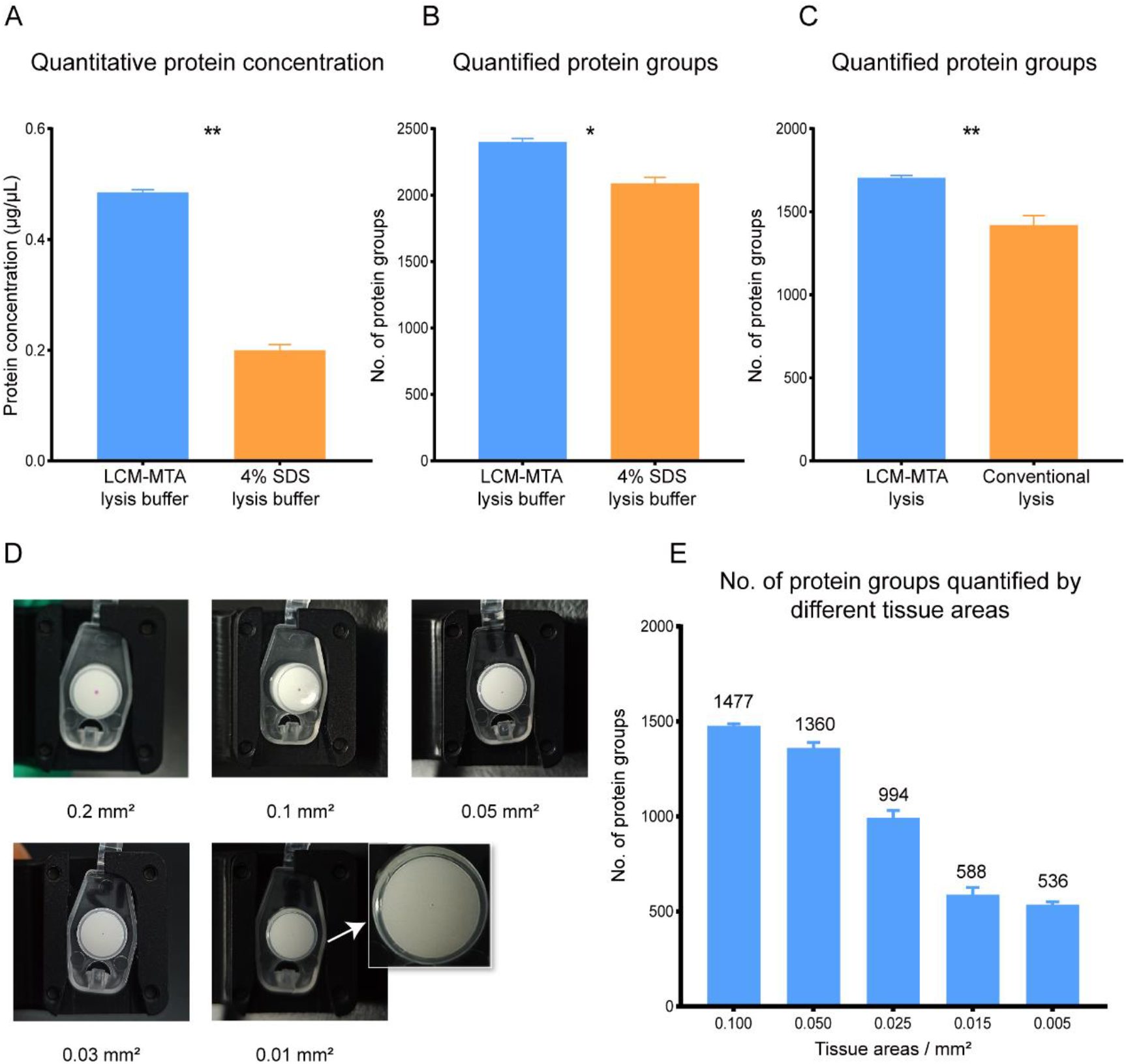
LCM-MTA can improve the depth of protein identification in trace FFPE tissues and reduce the starting tissues to 15 cells. **A**. Comparison of extracted protein concentration between LCM-MTA lysis buffer and 4% SDS lysis buffer. **B**. Comparison of the number of identified proteins between LCM-MTA lysis buffer and 4% SDS lysis buffer. **C**. Comparison of the number of identified proteins between LCM-MTA lysis and conventional lysis. Data in the barplot are presented as mean ± SEM. Statistical differences were calculated by two-tailed student’ s t-test. ns: Statistically not significant; *p < 0.05; **p < 0.01; ***p < 0.001. **D**. Images of trace tissue on adhesive caps of various areas obtained by LCM. **E**. The plot of the number of proteins identified from trace tissues of various areas. Trace tissue samples were prepared as peptide samples with LCM-MTA and one-half of the peptide samples were analyzed by LC-MS/MS. Data in the barplot are presented as mean ± SEM.

### LCM-MTA can efficiently extract membrane proteins and matrisome proteins

Membrane proteins play important roles in cellular biological processes, such as bioenergy transport, cell adhesion, and signal transduction. However, one of the challenges in studying membrane proteins using MS-based proteomics is their hydrophobicity, low solubility, and difficulty in extraction^22^. Here, we performed Gene Ontology (GO) cellular component analysis of proteomics data from colorectal cancer (CRC) and placental tissues. Compared with the conventional filter assisted sample preparation (FASP)^23^ sample preparation methods, LCM-MTA can extract more membrane proteins (Figure 3A). Matrisome is a complex network of cross-linked proteins that are important regulators of cell proliferation, differentiation, and migration. Dysregulation of matrisome is associated with numerous pathological conditions^24,25^. However, matrisome proteins have extensive covalent crosslinking and high insolubility compared to most intracellular proteins, which leads to isolation and extraction of matrisome proteins as a challenge for proteomic analysis^26,27^. In this work, based on proteomic data in DDA mode from CT-P and CT-S tissues, we found that LCM-MTA was able to obtain a higher proportion of matrisome proteins compared to FASP digestion^23^, which was similar to the proportion of matrisome proteins obtained by Naba et al^28^ using enrichment methods (Figure 3B). Comparison of matrisome composition obtained by LCM-MTA and other methods revealed that core matrisome accounted for a similar proportion (Figure 3C). We also observed that ECM glycoproteins, collagens, ECM regulators, ECM-affiliated proteins, proteoglycans, and ECM-associated secreted factors accounted for similar proportions of matrisome among the different methods (Figure 3D). In addition, similar results were obtained in data-independent acquisition (DIA) mode proteomic data of CT-P and CT-S tissues, suggesting that LCM-MTA enriched matrisome proteins were ubiquitous and unbiased (Figure 3E-G). It suggests that LCM-MTA can efficiently extract membrane proteins and matrisome proteins, which increases the integrity of SRP.

**Figure 3.**
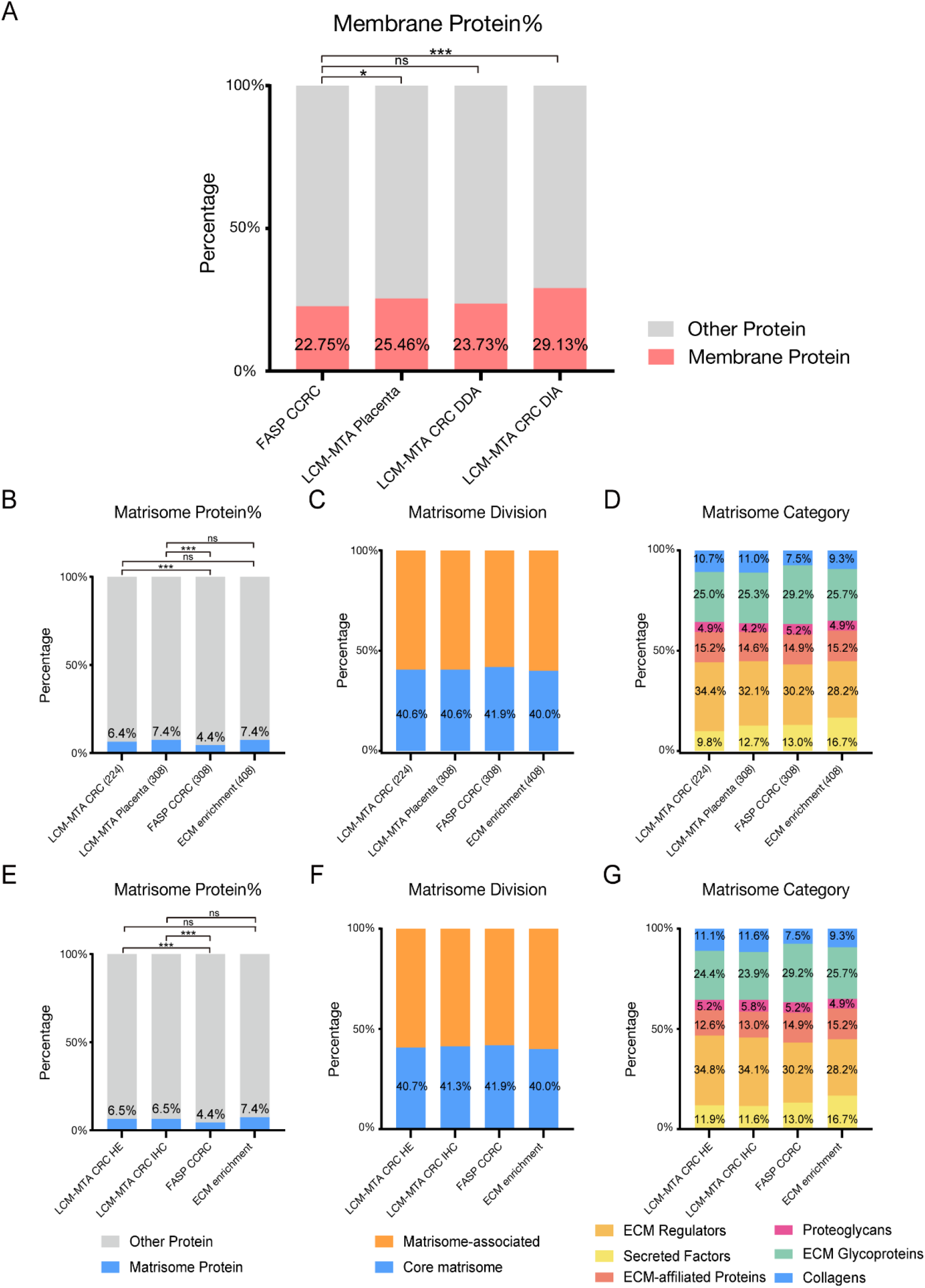
LCM-MTA can efficiently extract membrane proteins and matrisome proteins. **A**. The proportion of membrane proteins obtained during sample preparation using LCM-MTA and FASP in CRC and placenta tissues. Statistical differences were calculated by Fisher’s exact test. ns: Statistically not significant; *p < 0.05; **p < 0.01; ***p < 0.001. **B**. The proportion of matrisome proteins obtained during sample preparation using LCM-MTA, FASP, and enrichment methods in CRC and placenta tissues from DDA mode. Statistical analysis was the same as A above. **C**. The distribution of core matrisome and matrisome-associated proteins in matrisome protein obtained in B. **D**. The distribution of ECM regulators, secreted factors, ECM-affiliated proteins, proteoglycans, ECM glycoproteins, and collagens in matrisome protein obtained in B. The number in parenthesis indicate the amount of matrisome protein. **E**. The proportion of matrisome proteins obtained during sample preparation using LCM-MTA, FASP, and enrichment methods in CRC tissues from DIA mode. Statistical analysis was the same as A above. **F**. The distribution of core matrisome and matrisome-associated proteins in matrisome protein obtained in E. **G**. The distribution of ECM regulators, secreted factors, ECM-affiliated proteins, proteoglycans, ECM glycoproteins, and collagens in matrisome protein obtained in E.

### LCM-MTA has stability, universality, and extensibility

To illustrate the universality and extensibility of LCM-MTA, we compared the number of protein quantifications for different section thicknesses, different staining methods, and different LCM instruments in the DIA mode (Figure 4A-D). We compared the number of proteins identified by LCM-MTA in FFPE tissues with different staining methods and revealed that more proteins were identified in IHC stained tissues (Figure 4A-B), however the categories of overall proteins identified in each group were similar (Figure 4C), indicating the compatibility of LCM-MTA for FFPE tissues with different staining methods. In addition, For the same area of starting material, there was no significant difference in the number of proteins identified for FFPE tissue sections of different thicknesses (Figure 4B). When LCM was performed with different instruments, there was no significant difference in the number of proteins identified by LCM-MTA, indicating the compatibility of LCM-MTA for different LCM instruments (Figure 4D).

**Figure 4.**
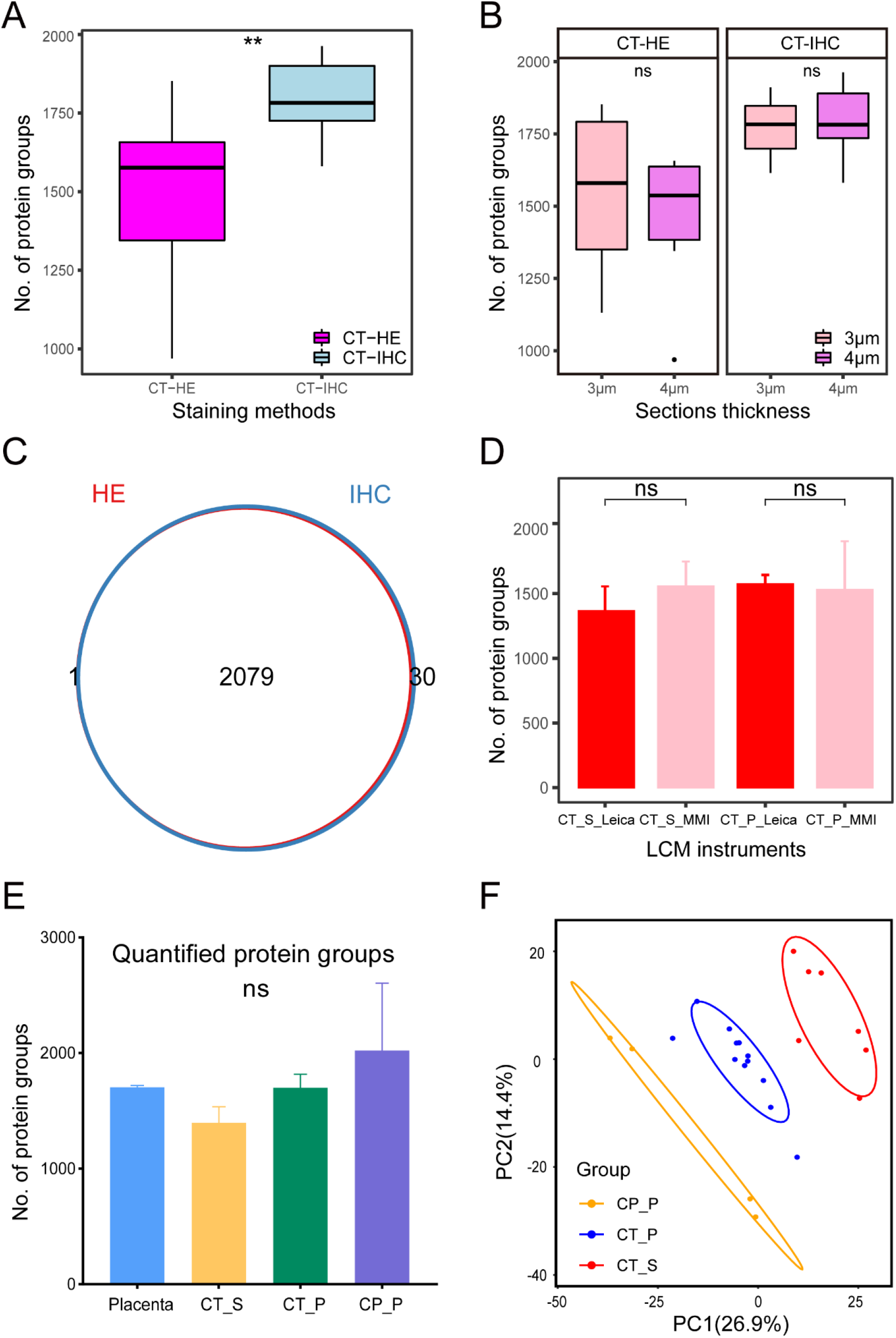
LCM-MTA has stability, universality, and extensibility. **A**. Comparison of the number of identified proteins between HE-stained tissues and IHC-stained tissues. **B**. Comparison of the number of identified proteins between 3-μm and 4-μm-thick sections. The line and box represent median and upper and lower quartiles, respectively. **C**. Venn diagram illustrates the overlap of identified proteins between HE-stained and IHC-stained tissues. **D**. Comparison of the number of proteins identified by MMI and Leica in DDA and DIA modes, respectively. Data in the barplot are presented as mean ± SEM. Statistical differences were calculated by two-tailed student’ s t-test. ns: Statistically not significant; *p < 0.05; **p < 0.01; ***p < 0.001. **E**. Comparison of the number of identified proteins between placenta, CT-S tissue, CT-P tissue, and CP-P tissue from global DDA and DIA proteomic data. CT-S, CRC CT stroma; CT-P, CRC CT tumor parenchyma; CP-P, CRC CP parenchyma. **F**. Principal Component Analysis distinguished CT tumor parenchyma from CP parenchyma and CT stroma based on global proteomic data from DDA and DIA mode.

Moreover, we integrated and compared DDA and DIA mode-based proteomic data derived from different kinds of tissues. There was no significant difference in the number of proteins identified across tissue types, indicating the stability of SRP based on LCM-MTA (Figure 4E). We also integrated and analyzed proteomic data based on DDA and DIA for CRC tissues derived from different staining methods, different section thicknesses, and different LCM instruments. PCA distinguished CT tumor parenchyma from CP parenchyma and CT stroma based on global proteomic data (Figure 4F). And it also shows the compatibility of LCM-MTA for different modes of MS data acquisition, different staining methods, different section thicknesses, and different LCM instruments. All the above indicated that LCM-MTA has stability, universality, and extensibility for SRP analysis of trace tissue samples.

### Spatially resolved cell-type protein profiles were characterized in clinical colorectal cancer tissues using LCM-MTA

To verify the reliability of LCM-MTA in SRP studies, we performed LCM-MTA and MS analysis on clinical CRC FFPE tissues. We obtained CT tumor parenchyma (CT-P), CT stroma (CT-S), CP parenchyma (CP-P), and CP stroma (CP-S) from CRC FFPE tissue by LCM (Figure 5A). CP-S tissues were not included in the analysis due to the low data available. Subsequently, trace target tissues were subjected to LCM-MTA and MS analysis. Following integration of proteomics data in DDA and DIA patterns from CT-S, CT-P, and CP-P tissues, we performed GO enrichment analysis. Consistent with the above results, membrane proteins and matrisome proteins were significantly enriched in CRC tissues (Figure 5B). Subsequently, principal Component Analysis (PCA) of global proteomic data from DDA and DIA mode can significantly distinguish tumor parenchyma from stroma (Figure 5C). Compared with CT-S tissue, CT-P tissue was characterized by increased metabolism of RNA, RNA splicing, ribonucleoprotein complex biogenesis, and ribosome biogenesis. Downregulated proteins in CT-P tissue were enriched for focal adhesion and supramolecular fiber organization (Figure 5D). In addition, to further validate the reliability of LCM-MTA in SRP studies, we also gathered CRC CP tissues, and then collected CP-P tissue by LCM. PCA clearly distinguished CT-P tissue from CP-P based on global DDA and DIA proteomic data after LCM-MTA and MS analysis (Figure 5E). And we found that compared with CP-P tissue, upregulated proteins in CT-P tissue were enriched for mRNA metabolic process, organelle assembly, ribonucleoprotein complex biogenesis, and intracellular protein transport. And upregulated proteins in CP-P tissue were enriched for metabolism of RNA, translation, and metabolic related pathways (Figure 5F). All the above indicate that the necessity of developing LCM based proteomic detection technology for trace tissue samples and the reliability of LCM-MTA in SRP studies.

**Figure 5.**
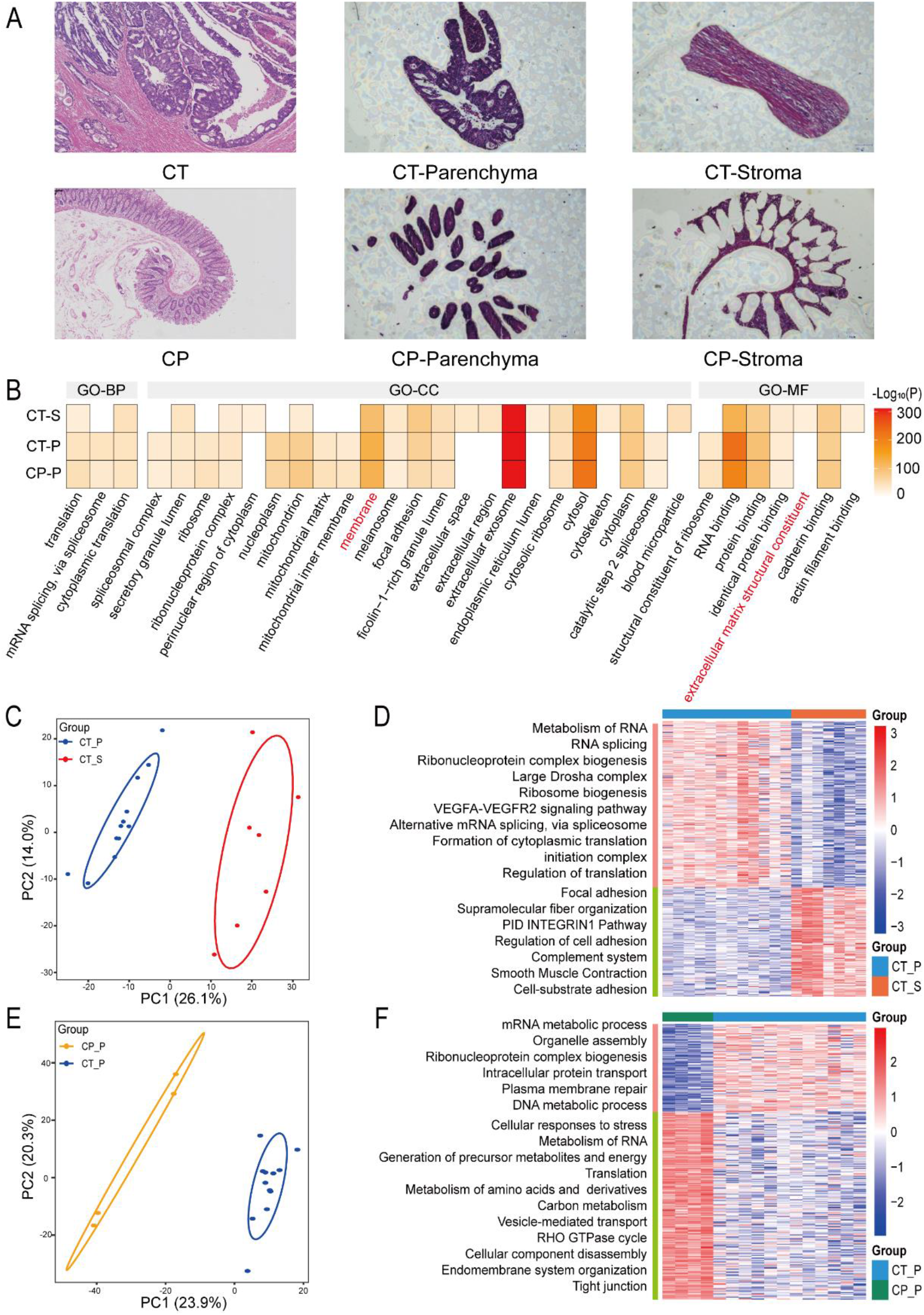
Spatially resolved cell-type protein profiles were characterized in CRC tissues using LCM-MTA. **A**. Images of parenchyma and stroma collected by LCM from CRC CT (top) and CP (bottom) tissues. **B**. Gene Ontology (GO) enrichment analysis of CRC global proteomic data from DDA and DIA mode. Only TOP 25 terms for each tissue type were shown in the plot. Red highlighted words indicate extracellular matrix structural constituent and membrane proteins. BP, biological process; CC, cellular component; MF, molecular function. **C**. Principal Component Analysis distinguished CRC CT tumor parenchyma from stroma based on global proteomic data from DDA and DIA mode. **D**. Normalized expression profiles of the differentially expressed proteins in CT P-S tissues and their enriched pathways. **E**. Principal Component Analysis distinguished CRC CT tumor parenchyma from CRC CP parenchyma based on global proteomic data from DDA and DIA mode. **F**. Normalized expression profiles of the differentially expressed proteins in CT-P and CP-P tissues and their enriched pathways. CT-P, CRC CT tumor parenchyma; CT-S, CRC CT stroma; CP-P, CRC CP parenchyma.

## Discussion

In this work, we provide an SRP sample preparation method suitable for trace FFPE tissues produced by LCM, termed LCM-MTA, which consists of five major stages. Briefly, trace samples of interest were obtained from FFPE tissue using LCM and collected on adhesive caps. Next, trace target tissues on adhesive caps were exposed to lysis buffer and subjected to lyse the tissues in situ. Subsequently, the lysate was centrifuged to collect proteins. Reducing agents and alkylating agents were added to the solution. And then beads and trypsin were added to the solution to digest proteins. Finally, digested peptides can be used for quantitative LC-MS/MS analysis.

FFPE is currently a standardized method for long-term stable preservation of clinical tissue specimens and provides a valuable resource for clinical research^29^. However, SRP analysis with FFPE tissues faces several challenges. The first challenge is that formaldehyde solution in FFPE tissues fixes the structure of the specimens by forming cross-linked proteins, which causes low efficiency of protein extraction during proteomics sample preparation^18^, especially for trace samples. Sigdel et al.^19^ reported a Near-Single-Cell proteomic study of kidney tissues, but the amount of protein extracted from FFPE sections was unsatisfactory. In this work, we present a method for lysis in situ that enables more efficient recovery of proteins compared to conventional lysis modality, thereby increasing the number of proteins identified. The second is that trace tissue samples are easily lost during proteomic sample preparation. Problematically, excessive target tissue area can drastically reduce spatial resolution, resulting in information homogenization in particular tissues or cell types. To improve spatial resolution, trace samples typically contain only a few dozen cells and contain proteins as low as sub-microgram levels. Herrera et al.^20^ reported a spatially resolved micro-proteomics technique in which they quantified more than 1,000 protein groups from 0.0125 mm^3^ (2.5 mm^2^ × 5 μm) human lung FFPE tissues, but the spatial resolution is not satisfactory. Our results show that more than 500 proteins can still be quantified at trace tissue size as small as 0.00004 mm^3^ (∼15 cells, 0.005 mm^2^ × 8 μm) based on DDA mode, which resolution is comparable to existing SRT^5,21^. And we will try to continue improving the spatial resolution based on DIA mode. And the third challenge is the detectability of MS for trace tissue samples, which is also one of the challenges faced by MS-based proteomics. Herein, we used a trapped ion mobility spectrometry coupled to time-of-flight mass spectrometer (timsTOF MS) combined with a high performance applied chromatographic system nanoElute® to improve the detectability of MS for trace tissues.

In addition, LCM-MTA has several other advantages. First, LCM-MTA can efficiently extract membrane proteins and matrisome proteins, which significantly increases the integrity of SRP. Membrane proteins are involved in various physiological and biochemical processes and are essential for cellular functions, including signal transduction and nutrient transport^30^. Membrane proteins also interact with cytoplasm, cytoskeleton, and other proteins to maintain cell or tissue homeostasis^31^. However, MS-based proteomics is challenging to characterize membrane protein profiles due to their hydrophobic, low solubility, and difficult extraction properties^32^. Therefore, scientists have developed strategies for enrichment, isolation, and proximity labelling techniques to characterize the protein profile of membrane proteins^31,33^. In this work we were surprised to find that LCM-MTA was able to extract more membrane proteins than the FASP sample preparation method. In addition to membrane proteins, we also found that LCM-MTA could extract more matrisome proteins. Matrisome is composed of collagen, glycoprotein, proteoglycan, and other components, which interact with surrounding cells to create a dynamic microenvironment and regulate cell and tissue homeostasis^34^. Matrisome proteins interacts with cells can regulate a variety of cellular functions, including proliferation, migration and differentiation, and dysregulation of matrisome is associated with the development and progression of various diseases such as cardiac diseases, neurodegenerative disorders, fibrosis, and cancers^35-39^. Problematically, matrisome proteins have extensive covalent crosslinking, high insolubility, and extensive post-translational modification compared to most intracellular proteins, which leads to isolation and extraction of matrisome proteins as a challenge for proteomic analysis^26,27^. Therefore, characterization of matrisome requires enrichment of matrisome proteins during sample preparation^25,28,40-42^. Surprisingly, LCM-MTA could extract more matrisome proteins than FASP^23^, and ECM glycoproteins, collagens, ECM regulators, ECM-affiliated proteins, proteoglycans, and ECM-associated secreted factors accounted for similar proportion of matrisome as other enrichment methods yielded. This will help us to pay more attention to the dynamic changes of matrisome proteins during the disease process in proteomic studies. Second, LCM-MTA has stability, universality, and extensibility. We have shown that LCM-MTA is compatible for different staining methods, section thicknesses, LCM instruments, and MS data acquisition modes, which means that LCM-MTA can be more widely used during practical research. Third, LCM-MTA is reliable in SRP studies. In this work, we demonstrate the reliability of LCM-MTA for SRP using clinical CRC FFPE tissues as an example. Compared with CT-S tissue, CT-P tissue was characterized by increased metabolism of RNA, RNA splicing, ribonucleoprotein complex biogenesis, and ribosome biogenesis. And compared with CP-P tissue, upregulated proteins in CT-P tissue were enriched for mRNA metabolic process, organelle assembly, ribonucleoprotein complex biogenesis, and intracellular protein transport. It also suggests apparent heterogeneity in clinical tissue bulk samples. Compared with bulk sample proteomics, SRP is able to focus on heterogeneity and spatial location information of clinical tissue samples^3^. More importantly, under classical histopathological guidance, information on the spatial distribution of proteins can reveal molecular features during tumorigenesis and progression, involving localized paracrine cytokine signaling and direct cell-cell contact^43^.Therefore, under the guidance of classical histopathology, in future work we can apply LCM-MTA to characterize the protein profile of specific regions of interest, such as tumor-stroma boundaries, to investigate tumor aggressiveness. In conclusion, it gives great confidence in our application of LCM-MTA to SRP studies, especially for special cell types or tissue types. Finally, LCM-MTA can easily implement high-throughput automation. By performing LCM-MTA on an automated liquid handling system, it can significantly accelerate and increase throughput for large-scale clinical samples, while reducing the variability of protein quantification and improving reproducibility.

In conclusion, LCM-MTA is a convenient, universal, and scalable method for SRP analysis of trace FFPE samples, which can be widely used in clinical and non-clinical research fields.

## Materials and methods

### Tissue samples acquisition and slides preparation

Human placenta tissues were collected from Shanghai First Maternity and Infant Hospital. Colorectal cancer (CRC) tissues were collected as previously described^23^. Briefly, we gathered primary tumors (CT) and paracarcinoma tissues (CP, 2-cm-away from the tumor edge, normal adjacent tissues) in this work. The collected tissue specimens were then prepared as FFPE tissues. Placental FFPE tissues were cut into 8-μm-thick sections and CRC FFPE tissues were cut into 3 μm, 4 μm, or 5 μm thick sections and mounted on MMI membrane slides, respectively, according to the experimental design. FFPE slides were dewaxed by treating them with xylene and alcohol, followed by HE or IHC staining, respectively.

### Laser capture microdissection

First, the prepared slides were loaded onto the stage of the MMI CellCut Laser Capture Microdissection System (Molecular Machines & Industries) or Leica Laser Capture Microdissection System (Leica LMD7). We then used a drawing tool on the computer screen to select regions of interest for microdissection. For 8-μm-thick sections of human placental tissue, we cut areas of approximately 0.01 mm^2^, 0.03 mm^2^, 0.05 mm^2^, 0.1 mm^2^, and 0.2 mm^2^, respectively, with two replicates for each group. In addition, to compare the protein extraction efficiency with different lysis buffers and different lysis modalities, we also collected placental tissues with an area of approximately 1 mm^2^ and 0.3 mm^2^, in two or three replicates per group, respectively. For 5-μm-thick sections of CRC CT and CP tissues, we cut parenchyma (CT-P, CP-P) and stroma (CT-S, CP-S) with an area of about 0.3 mm^2^, respectively. And for the 3-μm and 4-μm-thick sections of CRC CT tissues, we collected CT-P tissues and CT-S tissues with an area of approximately 0.3 mm^2^. For the MMI CellCut Laser Capture Microdissection System, we used the MMI adhesive caps to collect and store the target tissues on the adhesive caps, while for the Leica Laser Capture Microdissection System, we collected the target tissues in 96-well plates. Captured tissues were stored for later use.

### Protein extraction

To solubilize proteins from target tissues on the adhesive caps, we added lysis buffer of LCM-MTA directly to the caps. Samples were then incubated in situ in a water bath. After it had cooled to room temperature, the lysate was centrifuged and collected to the bottom of the tubes for later use. To compare the protein extraction efficiency of LCM-MTA lysis buffer and conventional SDS lysis buffer, the LCM-MTA lysis buffer or 4wt% SDS lysis buffer was added directly to the caps. Samples were incubated at room temperature, followed by centrifugation to collect samples to the bottom of tubes. Samples were then incubated in a water bath. Moreover, to compare the protein extraction efficiency of different lysis modalities, the LCM-MTA lysis buffer was added directly to the caps. The test group was subjected to water bath in situ as described above, while the control group was incubated at room temperature followed by centrifugation and then incubated in a water bath. Protein concentration was detected using tryptophan-based fluorescence quantification method^44^.

### Peptides preparation

We then added 0.5 M chloroacetamide (CAA) and 0.5 M Tris (2-carboxyethyl) phosphine (TCEP) to the protein mixture, following by incubation at 95 °C for 5 min. After it had cooled to room temperature, beads were added to protein samples, and then ethanol was added to make the final concentration of ethanol 50 wt%. Samples were shaken on a mixer with mixing at 1,000 r.p.m. The samples were placed on the magnetic rack and incubated, and then the supernatant was removed and discarded. Subsequently, wash the beads with 80% (vol/vol) ethanol, remove and discard the supernatant. Repeat this step once. And then wash the beads by adding 100% ethanol and remove and discard the supernatant as above. After the beads were air-dried, 100 mM Ammonium bicarbonate was added to the beads and then the samples were sonicated in a water bath sonicator at room temperature. Subsequently, 0.1 μg/μL trypsin (Thermo Fisher Scientific) in Ammonium bicarbonate was added to the samples, followed by incubation at 37 °C in a mixer at 1,000 r.p.m. mixing. Following digestion, the tubes were placed on the magnetic rack and incubated until the beads migrated to the wall. The supernatant was then transferred to the new tubes and 1% (vol/vol) formic acid was added to acidify the peptide mixture.

### LC-MS/MS analysis

Peptide samples obtained from the above steps was used for MS-based proteomic analysis. The LC-MS/MS analysis was performed on a trapped ion mobility spectrometry coupled to time-of-flight mass spectrometer (timsTOF Pro, Bruker) combined with a high performance applied chromatographic system nanoElute® (Bruker). Peptides were loaded on to an in-house packed column (75 μm × 250 mm; 1.9 μm ReproSil-Pur C18 beads, Dr. Maisch GmbH, Ammerbuch) which was heated to 60 °C, and separated with a 60-min gradient of 2% to 80% mobile phase B at a flow rate of 300 nL/min. The mobile phases A and B were 0.1% (v/v) formate/ddH_2_O and 0.1% (v/v) formate/acetonitrile, respectively. The mass spectrometer was performed with default parameters either in a data-dependent acquisition (DDA) or data-independent acquisition (DIA) parallel accumulation-serial fragmentation (PASEF) mode^45^.

### Database searching

DDAPASEF data were analyzed on PEAKS software (Version Online X) and searched against the UniProt/SwissProt human database (20,350 entries, May 25, 2020) which following the standard workflow. DIAPASEF data were analyzed on the Spectronaut™ (version 15.0) with the spectral libraries generated by Pulsar algorithm using both DDAPASEF and DIAPASEF files.

### Data analysis

Statistical significance tests, including student’s t-test, and Fisher’s exact test were performed using R v.4.1.1, as denoted in each analysis. Data in the barplot are presented as mean ± SEM, and in boxplot the line and box represent median and upper and lower quartiles, respectively. All statistical tests were two-sided, and statistical significance was considered when P value < 0.05. CP-S tissues were not included in the data analysis due to the low data available. The protein expression levels were normalized using the median centering methods and log2-transformed. ComBat functions in “sva” R package were applied to eliminate batch effect^46^. The Principal Component Analysis (PCA) was performed in R v.4.1.1. Differential proteomic analysis of CRC tissues was performed for proteins quantified in at least half of the samples. Protein expression differences were analyzed using the student’ s t-test if protein data were normally distributed, otherwise Wilcoxon rank-sum test. Differentially expressed proteins were defined as those showing fold change > 3/2 or < 2/3 and P value (the P values were adjusted using the Benjamini-Hochberg FDR correction) < 0.05. Matrisome data were derived from MatrisomeDB database^47^. Functional enrichment analysis was performed using Metascape^48^ and DAVID^49^ with default parameters.

## Acknowledgements

We thank all members of our laboratories for helpful discussions. This work was supported by the National Natural Science Foundation of China (NSFC) (grants 82030099, 30700397), the Shanghai Municipal Science and Technology Commission “Science and Technology Innovation Action Plan” technical standard project (21DZ2201700), and the innovative research team of high-level local universities in Shanghai.

## Author contributions

Conceptualization, H.Q.Z., Z.W., L.C. and W.H.; Methodology & Investigation, L.C., G.L., L.X.M., L.Z.Y, W.Q.Q., Z.K., Y.G.Y. and D.C.Q; Writing – Original Draft, L.C., L.X.M, G.L. and L.Z.Y.; Writing – Review & Editing, W.Q.Q., Z.K., Y.G.Y., D.C.Q., L.J.Q., Z.B.P. and Z.H.P; Supervision, H.Q.Z., Z.W., L.C. and W.H.

